# A physics-based energy function allows the computational redesign of a PDZ domain

**DOI:** 10.1101/790667

**Authors:** Vaitea Opuu, Young Joo Sun, Titus Hou, Nicolas Panel, Ernesto J. Fuentes, Thomas Simonson

## Abstract

A powerful approach to understand protein structure and evolution is to perform computer simulations that mimic aspects of evolution. In particular, structure-based computational protein design (CPD) can address the inverse folding problem, exploring a large space of amino acid sequences and selecting ones predicted to adopt a given fold. Previously, CPD has been used to entirely redesign several proteins: all or most of the protein sequence was allowed to mutate freely; among sampled sequences, those with low computed folding energy were selected, and a few percent of them did indeed adopt the correct fold. Those studies used an energy function that was partly or largely knowledge-based, with several empirical terms. Here, we show that a PDZ domain can be entirely redesigned using a “physics-based” energy function that combines standard molecular mechanics and a recent, continuum electrostatic solvent model. Many thousands of sequences were generated by Monte Carlo simulation. Among the lowest-energy sequences, three were chosen for experimental testing. All three could be overexpressed and had native-like circular dichroism and 1D-NMR spectra. Two exhibited an increase in their thermal denaturation curves when a peptide ligand was present, indicating binding and suggesting correctly folded proteins. Evidently, the physical principles that govern molecular mechanics and continuum electrostatics are sufficient to perform whole-protein redesign. This is encouraging, since these methods provide physical insights, can be systematically improved, and are transferable to other biopolymers and ligands of medical or technological interest.

Table of Contents Graphic

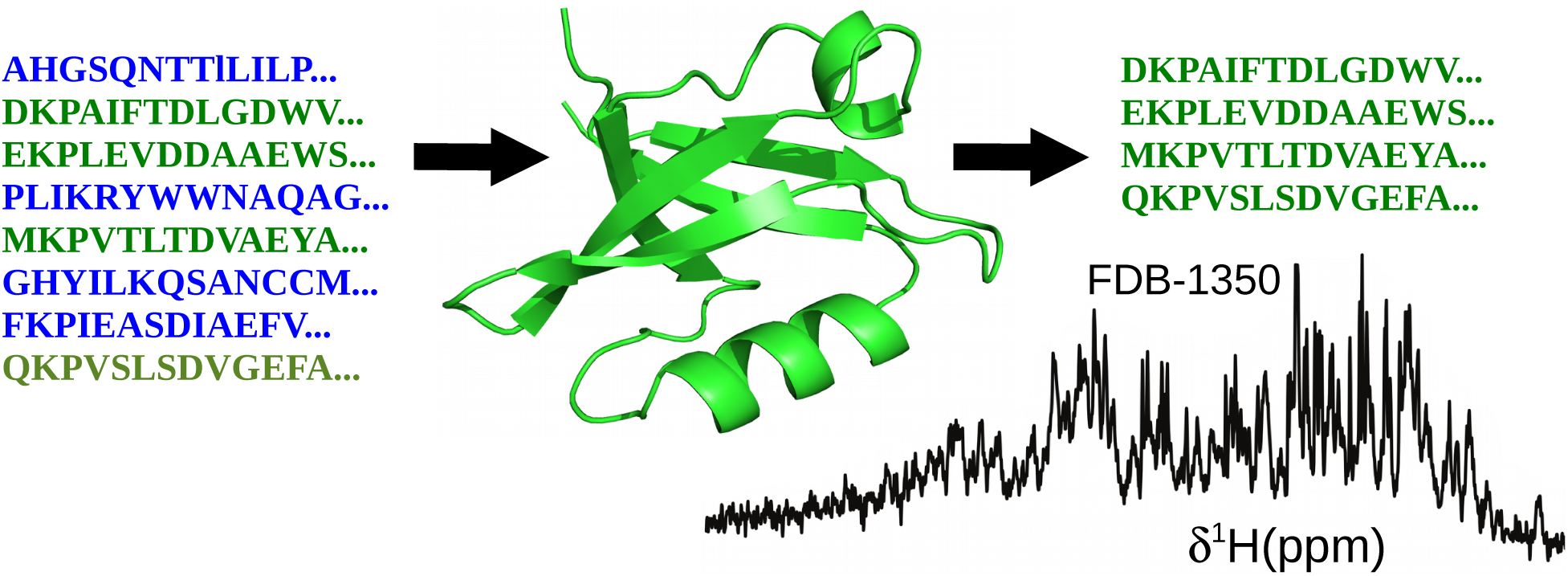

## Introduction

Protein sequences have been selected by millions of years of evolution to fold into specific 3D structures, stabilized by a subtle balance of interactions involving protein and solvent.^1–3^ In contrast, random polymers of amino acids are very unlikely to adopt a specific, stable, folded structure, ^4–6^ and exhibit instead a more disordered structure. ^7^ A powerful approach to understand the evolution of proteins and the physical origins of folding is to perform simulations that mimic evolution in a computer. This can be done with computational protein design (CPD), which explores a large set of amino acid sequences and selects ones that are predicted to adopt a given fold. ^8–12^ A typical simulation imposes a specific geometry for the protein backbone, corresponding to the experimental conformation of a natural protein. Amino acid side chains are mutated randomly, for example through a Monte Carlo procedure. Variants that have a favorable predicted folding free energy are saved. The energy of the folded state is predicted with an energy function that can be physics-based or knowledge-based. ^13–16^ The unfolded state is usually described by a simple energy function that depends on the protein sequence but does not involve a detailed structural model. The protein is considered “completely redesigned” if most or all of the protein side chains are allowed to mutate during the simulation. Indeed, the final, predicted sequences will then have low sequence identity to the natural protein whose backbone was used as a starting point.

The successful redesign of several complete proteins was first reported in 2003. ^9, 17^ The calculations were based on the Rosetta energy function, which contains several empirical terms parameterized specifically for protein design. Therefore, it can be considered to be at least partly knowledge-based. Several other successes were obtained^18, 19^ with updated versions of the Rosetta energy function, ^16^ including a recent large-scale study where 15000 miniproteins (40–43 amino acids) were redesigned. ^20^ 6% of the 15000 designs were shown to be successful; i.e., the designed miniproteins folded into the correct 3D conformation. The others either could not be overexpressed and purified, or did not fold as predicted. In addition to Rosetta, other knowledge-based energy functions were used to successfully redesign several proteins. ^21, 22^

Energy functions for the folded state can also be taken from molecular mechanics. ^23, 24^ There are then only two energy terms for nonbonded interactions between protein atoms, which correspond to the elementary Coulomb and Lennard-Jones effects. Their parameterization relies mainly on fitting quantum chemical calculations performed on small model compounds in the gas phase. The solvent is described implicitly, using varying levels of approximation. ^25^ The most rigorous model used so far for CPD is a dielectric continuum model. ^14, 26^ This requires solving a differential equation, which is technically impractical in a protein design framework. Therefore, a Generalized Born (GB) approximation is more common. GB contains much of the same physics but provides a simpler, analytical energy expression. ^25, 27^ GB models have been studied extensively in the context of protein design but also molecular dynamics, free energy simulations, acid/base calculations, ligand binding and protein folding. ^25, 28–32^ They reproduce the behavior of the dielectric continuum model rather accurately. Therefore, an energy function that combines molecular mechanics for the protein with a Generalized Born solvent can be considered “physics-based”, even though it is not entirely constructed from first principles. A molecular mechanics energy, combined with a very simple solvent model, was used to computationally design two artificial proteins that each consisted of a four-helix bundle, where an elementary unit of 34 amino acids was replicated four times. ^33, 34^ However, until now, there had not been a complete redesign of a natural protein using a physics-based energy function.

Here, we report the successful use of a physics-based energy function to completely redesign a PDZ domain of 83 amino acids. PDZ domains (“Postsynaptic density-95/Discs large/Zonula occludens-1”) are globular domains that establish protein-protein interaction networks.^35–37^ They interact specifically with target proteins, usually by recognizing a few amino acids at the target C-terminus. They have been extensively studied and used to elucidate principles of protein evolution and folding. ^38–40^ Our design started from the PDZ domain of the Calcium/calmodulin-dependent serine kinase (CASK) protein. It used the backbone conformation from an X-ray structure reported here. Positions occupied by glycine (seven) or proline (two) were not allowed to mutate. 13 positions involved in peptide binding also kept their wildtype identity. All 61 of the other side chains (73.5% of the sequence) were allowed to mutate freely into any amino acid type except Gly or Pro, for a total of 3.7 10^76^ possible sequences. The energy function combined the Amber ff99SB molecular mechanics force field ^41^ and a GB solvent model. ^27, 42^ Computations were done with the Proteus software. ^43, 44^ Three designs were tested experimentally and were all shown to fold. For two, binding to one or two peptides that are known CASK ligands was demonstrated. Evidently, the physical principles that govern molecular mechanics and continuum electrostatics are sufficient to allow large-scale computational protein design. This is encouraging, since these methods give physical insights, can be systematically improved, and are transferable to nucleic acids, sugars, noncanonical amino acids, biological cofactors, and many ligands of therapeutic or biotechnological interest.

## Materials and methods

### Computational design methods

#### Energy function for the folded state

We used the following effective energy function for the folded state:

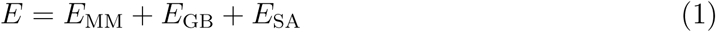

*E*_MM_ is the protein internal energy, taken from the Amber ff99SB molecular mechanics (MM) energy function. ^41^ *E*_GB_ is a Generalized Born (GB) implicit solvent contribution: ^27, 45^

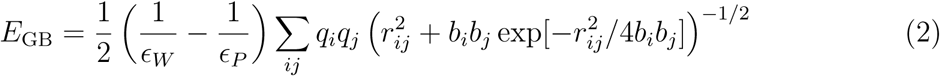

Here, *∊_W_* and *∊_P_* are the solvent and protein dielectric constants (80 and 4, respectively); *r_ij_* is the distance between atoms *i, j* and *b_i_* is the “solvation radius” of atom *i*.^27, 42^ The dependency of the *b_i_* on the protein conformation corresponds to a GB variant we call GB/HCT (for “Hawkins-Cramer-Truhlar”). ^27, 42^ For some of the design calculations, an additional “Native Environment Approximation”, or NEA was used for efficiency,^45, 46^ where the solvation radius *b_i_* of each particular group (backbone, sidechain or ligand) was computed ahead of time, with the rest of the system having its native sequence and conformation. ^46, 47^ For the other designs, we computed the solvation radii on the fly during the MC simulation, using a very fast implementation called “Fluctuating Dielectric Boundary,” or FDB^47^ that uses lookup tables.

The last term in Eq. (1) is a surface area term:

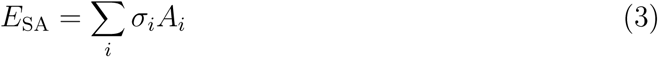

*A_i_* is the exposed solvent accessible surface area of atom *i*; *σ_i_* is a parameter that reflects each atom’s preference to be exposed or hidden from solvent. The solute atoms were divided into four groups with specific *σ_i_* values. The values were −60 (nonpolar), 30 (aromatic), −120 (polar), and −110 (ionic) cal/mol/Å^2^. The coefficient for hydrogens was zero. Surface areas were computed by the Lee and Richards algorithm, ^48^ implemented in the Proteus software, ^43^ using a 1.5 Å probe radius. To avoid overcounting buried surface, a scaling factor of 0.65 was applied to the contact areas involving at least one buried side chain. ^42, 45^

#### The unfolded state model

For a particular sequence *S*, the unfolded state energy has the form:

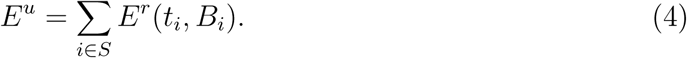

The sum is over all amino acids; *t_i_* represents the side chain type at position *i*; *B_i_* represents the buried or exposed character of position *i* in the folded state. The quantities *E^r^*(*t, B*) *≡ E^r^_t_* are referred to as “reference energies”; they can be thought of as effective chemical potentials of each amino acid type. Their values were chosen to maximize the likelihood of a set of experimental PDZ sequences. In practice, this means that a Monte Carlo simulation should give amino acid frequencies that match those in the experimental sequences. ^49^ We assigned different values to buried and exposed positions, because we assume residual structure is present in the unfolded state, so that amino acids partly retain their buried/exposed character. To define the target amino acid frequencies for likelihood maximization, we used a set of PDZ sequences collected earlier. ^49^

#### Structural model and energy matrix

For CASK, we used a new X-ray structure of the apo PDB domain, reported here (PDB entry 6NH9). To carry out the Monte Carlo simulations, an energy matrix was computed using procedures described previously.^49^ Briefly, for each pair of amino acid side chains, the interaction energy was computed after 15 steps of energy minimization, with the backbone held fixed and only the interactions of the pair with each other and the backbone included. ^50^ Side chain rotamers were described by the Tuffery library,^51^ expanded to include additional hydrogen orientations for OH and SH groups. ^45^ The energies were stored in an energy table, or “matrix” for use during MC.

#### Monte Carlo simulations

Sequence design was performed by running long Monte Carlo (MC) simulations where 61 out of 83 positions could mutate freely: all but 7 Gly, 2 Pro and 13 positions that are directly involved in binding the peptide ligand. Non-mutating positions could explore different rotamers. The MC simulations used one- and two-position moves, where either rotamers, amino acid types, or both changed. For two-position moves, the second position was near the first in space. Sampling was enhanced by Replica Exchange Monte Carlo (REMC), where eight MC simulations (“replicas”) were run in parallel, at different temperatures. ^52^ Periodic swaps were attempted between the conformations of two replicas *i*, *j* (adjacent in temperature), subject to a Metropolis acceptance test.^52^ Thermal energies ranged from 0.125 to 3 kcal/mol. Simulations were done with the Proteus software. ^46, 52^

#### Sequence characterization

Designed sequences were compared to the Pfam alignment for the PDZ family, using the Blosum40 scoring matrix and a gap penalty of −6. Designed sequences were also submitted to the Superfamily library of Hidden Markov Models, ^53, 54^ which attempts to classify sequences according to the Structural Classification Of Proteins, or SCOP.^55^ The isoelectric point of each sequence was estimated by assuming each titratable side chain had its standard pK*_a_* value.

#### Molecular dynamics simulations

Wildtype CASK and six sequences designed with Proteus were subjected to MD simulations with explicit solvent and no peptide ligand. The starting structures were taken from the MC trajectory or the crystal structure and slightly minimized with harmonic restraints to maintain the backbone geometry. Each protein was immersed in a solvent box using the CHARMM GUI.^56, 57^ The boxes had a truncated octahedral shape. The minimum distance between protein atoms and the box edge was 15 Å. The final models included about 11,000 water molecules. A few sodium or chloride ions were included to ensure overall electroneutrality. The protonation states of histidines were assigned to be neutral, based on visual inspection. MD was performed with periodic boundary conditions, at room temperature and pressure, using Langevin dynamics with a Langevin Piston NośeHoover barostat.^58, 59^ Long-range electrostatic interactions were treated with a Particle Mesh Ewald approach. ^60^ The Amber ff14SB force field and the TIP3P model^61^ were used for the protein and water, respectively. Simulations were run for one microsecond, using the Charmm and NAMD programs. ^57, 62^

#### Protein expression and purification

The codon optimized gene of the human CASK PDZ domain (residues 487–572) was chemically synthesized (GenScript Inc., Piscataway, NJ) and ligated into the pET28a vector (Novagen). The DNA sequence of the pET28a-CASK PDZ vector was verified by automated DNA sequencing (University of Iowa, DNA Facility). Protein expression was conducted in BL21(DE3) (Invitrogen) *E. coli* cells. Typically, *E. coli* cells were grown at 37*^◦^*C in Luria-Bertani (LB) medium supplemented with kanamycin (15 *µ*g/mL) under vigorous agitation until an absorbance at 600 nm wavelength (A600) reached 0.6-0.8. Cultures were subsequently cooled to 18*^◦^*C and protein expression was induced by the addition of isopropyl 1-thio-*β*-d-galactopyranoside (IPTG) to 1 mM final concentration. Induced cells were incubated for an additional 1618 hrs at 18*^◦^*C. and harvested by centrifugation. The CASK PDZ domain was purified by cation exchange (SP media, GE-Healthcare) and size-exclusion chromatography (GE-Healthcare). The Superdex 75 (S75) size-exclusion chromatography was performed with desired final buffer (20 mM phosphate, pH 6.8, 50 mM NaCl, and 0.5 mM EDTA). The final yield was 50 mg of CASK PDZ protein ¿98% pure as judged by SDS-PAGE from 1 L of culture. Samples were used immediately or lyophilized and stored at −80*^◦^*C. The Tiam1 PDZ domain was purified as previously published.^63^

The genes encoding the Proteus PDZ designs were codon-optimized for *Escherichia coli* expression and chemically synthesized by GenScript Inc. (Piscataway, NJ). The genes were cloned into a modified pET21a vector (Novagen) that contains a His_6_-tag and Tobacco etch virus protease cleavage site at the 5′-end of the multiple cloning site. The nucleotide coding sequence of the pET21a-PDZ vector was verified by automated DNA sequencing (University of Iowa, DNA Facility). Protein expression was conducted in BL21(DE3) (Invitrogen) *E. coli cells*. Typically, cells were grown at 37*^◦^*C in Luria-Bertani medium supplemented with ampicillin (100 *µ*g/mL) under vigorous agitation until an A600 of 0.6-0.8 was reached. Cultures were subsequently cooled to 18*^◦^*C and protein expression was induced by the addition of ITPG to 1 mM final concentration. Induced cells were incubated for an additional 16-18 hrs at 18*^◦^*C and harvested by centrifugation. Proteins were initially purified by nickel-chelate chromatography (GE-Healthcare). The proteins were further purified by size-exclusion chromatography (Superdex 75, GE Healthcare) using a buffer containing 20 mM phosphate, pH 6.8, 50 mM NaCl, and 0.5 mM EDTA. Samples were used immediately.

### Crystal structure of the wildtype apo CASK PDZ domain

A crystal structure of the apo CASK PDZ domain was determined in this work. High-throughput hanging-drop, vapor-diffusion screens using a Mosquito drop setter (TTP LabTech) were used to determine the crystallization conditions. The CASK PDZ domain was prepared in 20 mM Tris pH 7.5 and 50 mM NaCl. 200 nL of precipitant and PDZ domain (10-30 mg/mL) was used for each screening condition. Initial screening for diffracting crystals was done with an in-house Rigaku RAXIS-IV rotating anode X-ray source. Collection of full X-ray diffraction datasets for structure determination was done at beamline 4.2.2 at the Advanced Light Source (Berkeley, CA). Proper space group handedness was verified by analysis of the electron density.

XDS was used for indexing, integration, and scaling of the diffraction data,^64, 65^ to 2.0 Å resolution. XSCALE was used to merge multiple datasets. We used PHASER and previously-determined PDZ structures for initial phasing. ^66^ We used Refmac^67, 68^ for the early stages of refinement and PHENIX^69, 70^ for the final refinement. Refinement statistics are given in Supplementary Material (Table S1). Manual model building was done based on visualized electron density in Coot.^71^ 4.6% of the reflections were randomly selected to be excluded from the refinement and used to calculate R_free_ values. Alignment of structures and generation of figures were done with PyMOL (Schrodinger, LLC, The PyMOL Molecular Graphics System).

**Table 1:** DSF for wildtype CASK and three Proteus designs

| protein<sup>a</sup> | $T_{1/2}$ (°C) and $\delta T_{1/2} = T_{1/2}^{\text{apo}} - T_{1/2}$ (in parentheses) | | | | binding<sup>b</sup> |
| --- | --- | --- | --- | --- | --- |
|  | Apo | SDC1 | Caspr4 | NRXN |  |
| CASK PDZ | $57.2 \pm 0.2$ | $58.4 \pm 0.1$<br>(+1.2) | $58.7 \pm 0.1$<br>(+1.5) | $58.1 \pm 0.2$<br>(+0.9) | **Sdc1, Caspr4**<br>NRXN |
| FDB-1350 | $49.8 \pm 0.4$ | $50.7 \pm 0.2$<br>(+0.9) | $50.4 \pm 0.4$<br>(+0.6) | $51.3 \pm 0.2$<br>(+1.5) | Sdc1<br>**NRXN** |
| FDB-1669 | $49.1 \pm 0.1$ | $49.6 \pm 0.1$<br>(+0.4) | $49.5 \pm 0.0$<br>(+0.4) | $50.1 \pm 0.1$<br>(+1.0) | **NRXN** |
| FDB-1555 | $49.9 \pm 0.2$ | $50.2 \pm 0.1$<br>(+0.3) | $50.3 \pm 0.1$<br>(+0.5) | $50.5 \pm 0.6$<br>(+0.6) | — |

<sup>a</sup>Protein concentration was $\sim 25 \mu\text{M}$ (about 0.25 mg/ml). <sup>b</sup>When $\delta T_{1/2}$ was larger than sum of the standard deviation of apo and each peptide, we considered the peptides to have a significant change in $T_{1/2}$ , indicating binding to the PDZ domain. ± indicates standard deviation of three biological replicates.

### Biophysical characterization of designed proteins

#### Synthetic peptides

All peptides were chemically synthesized by GenScript Inc. (Piscataway, NJ) and were *>*95% pure as judged by analytical HPLC and mass spectrometry. Peptides were dansylated at the N-terminus and had a free carboxyl at the C-terminus. The peptides used in this study were derived from the following proteins: Neurexin (residues 1,470-1,477: NKDKEYYV*_COOH_*), Caspr4 (residues 1,301-1,308: ENQKEYFF*_COOH_*) and Syndecan1 (residues 303310: TKQEEFYA*_COOH_*).

#### Circular dichroism

Circular dichroism signals were measured using a Jasco J-815 circular dichroism spectropolarimeter. The concentration of each protein ranged from 10 to 20 *µ*M. All proteins were in a buffer composed of 20 mM phosphate, pH 6.8, 50 mM NaCl, and 0.5 mM EDTA. Spectra were taken from the 190 nm to 260 nm wavelength window with a 1 nm data interval at 25*^◦^*C. Data integration time was 2 seconds and the scanning speed was 100 nm/min.

#### NMR

NMR experiments were carried out at 298 K (calibrated with methanol) on Bruker Avance II 800 MHz (equipped with a CryoProbe), Bruker Avance II 500 MHz, and Varian 600 MHz spectrometers (equipped with room temperature probes). All protein samples were prepared in 20 mM phosphate, pH 6.8, 50 mM NaCl, 0.5 mM EDTA, and 10% (v/v) D_2_O with a concentration of 14 *µ*M to 22 *µ*M.

#### Differential scanning fluorimetry

Standard methodology was used for differential scanning fluorimetry (DSF).^72, 73^ Briefly, DSF was performed using 96-well PCR plates and the Sypro Orange (Thermo Fisher) dye. Each well in the PCR plate had a 20 *µ*L final volume containing 0.25 mg/mL of protein, 300 *µ*M of peptide, and 5x Sypro Orange final concentration (from a 5000x stock) in a buffer containing 20 mM phosphate, pH 6.8, 50 mM NaCl, and 0.5 mM EDTA. The DSF assays were performed using a Bio-Rad CFX96 real-time polymerase chain reaction instrument equipped to read 96-well plates. The protein of interest was thermally denatured from 5*^◦^*C to 95*^◦^*C at a ramp rate of 1*^◦^*C/min. The protein melting/unfolding curves were generated by monitoring changes in Sypro Orange fluorescence (at 610 nm wavelength). Raw fluorescence data were analyzed using DMAN, and the first derivative value from the denaturation data was used to determine the apparent melting temperature ^74^ (T_1*/*2_). Each peptide was assayed in triplicate. A 96-well plate containing no peptide was assayed to determine the apparent T_1*/*2_ of each PDZ domain in the absence of any peptide. A shift of more than 1*^◦^*C in T_1*/*2_ indicates binding (based on SEM).

## Results

Protein design simulations were performed using the CASK backbone conformation, shown in Fig. 1. 61 out of 83 residues were allowed to mutate into all types except Gly and Pro, for a total set of 18^61^ = 3.7 10^76^ possible sequences. This set was explored using Replica Exchange Monte Carlo, without any bias towards natural sequences or any limit on the number of mutations. The 2,000 sequences with the lowest folding energies were retained for analysis. Below, we describe the computational characterization of the designed sequences and the selection of three representative sequences for experimental characterization.

**Figure 1:**
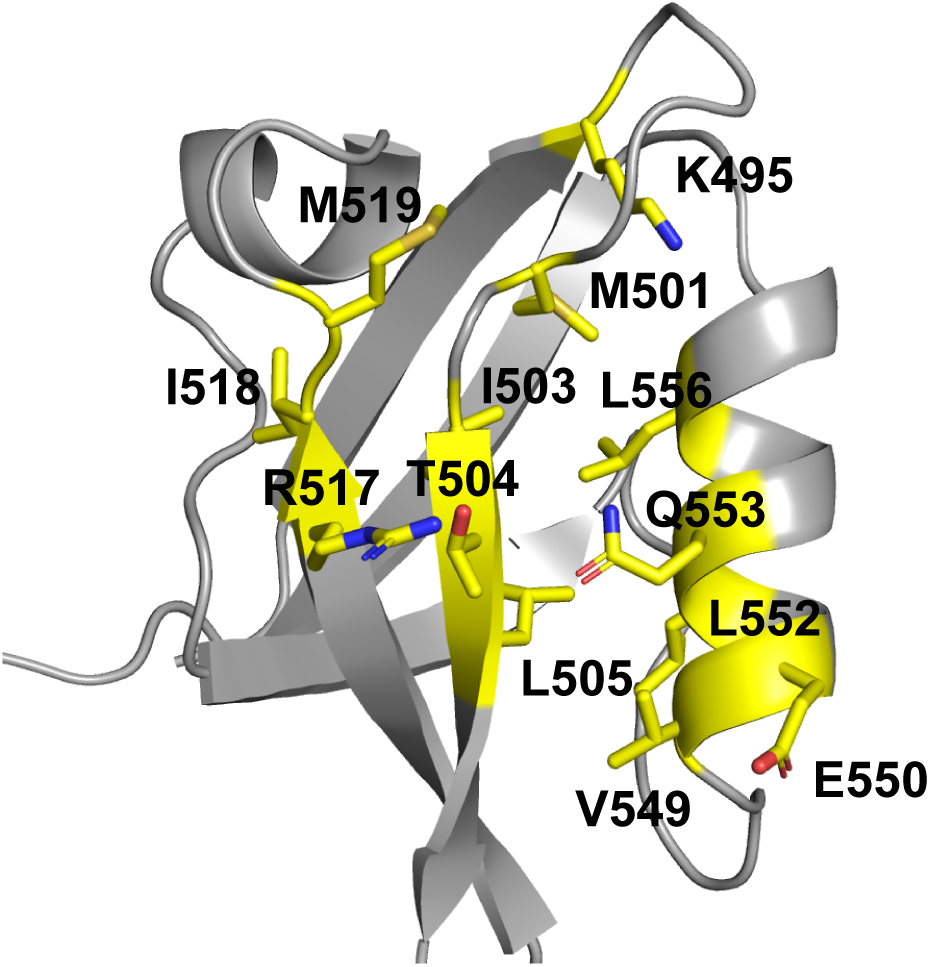
CASK 3D structure. Highlighted in yellow are the 13 amino acid positions whose identity was kept fixed in the Proteus designs.

### Computational characterization and sequence selection

The top 2,000 sequences spanned a folding energy interval of 1.5 kcal/mol. Since negative design was not included in our sequence generation, there is a possibility that some sequences could fold into a non-PDZ structure. Therefore, they were analyzed by the Superfamily fold recognition tool, ^54, 75^ which assigns sequences to SCOP structural families. All 2,000 Proteus sequences were assigned by Superfamily to the PDZ fold; none were predicted to adopt any other fold in SCOP. We next computed the Blosum40 similarity scores between the designed sequences and natural sequences from the Pfam database. Two histograms of scores are shown in Fig. 2: one for the whole protein and one for the protein core (15 positions). The scores of the sequences designed with Proteus (with 61 out of 83 positions allowed to mutate) are high, and comparable to the scores of natural PDZ domains. The peaks in the Proteus histograms are narrow, indicating that the 2,000 lowest-energy sequences are similar to each other.

**Figure 2:**
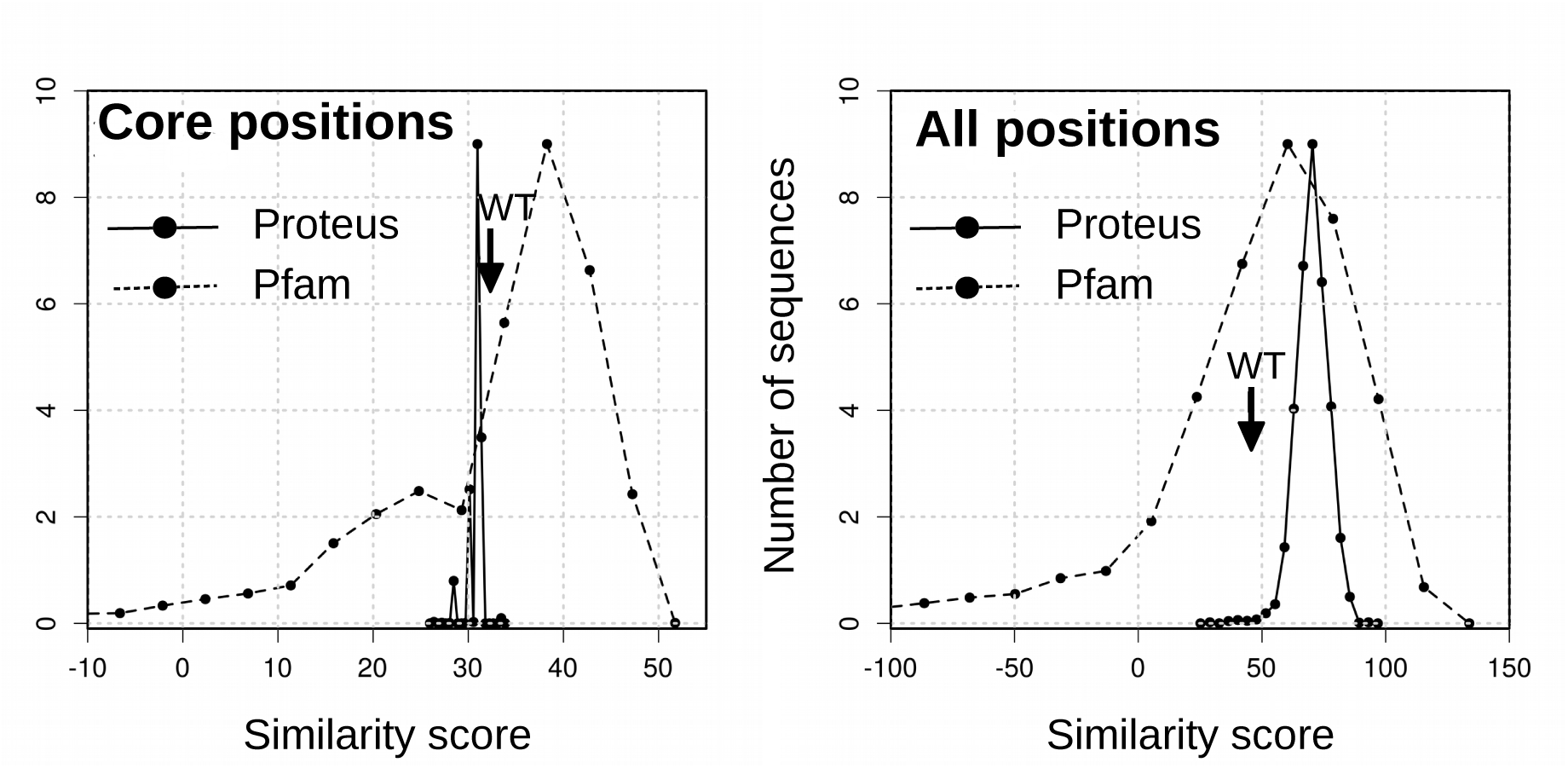
Blosum similarity scores compared to natural Pfam sequences. Black line: histogram of scores for the top 2000 Proteus sequences, considering only core positions (left) or all positions (right). Dashed line: scores for the Pfam sequences themselves. WT score is indicated by an arrow.

To narrow down the number of sequences, we excluded those with isoelectric points estimated to be close to the physiological pH, between 6.5 and 8.5, which might be subject to aggregation and difficult to express. This reduced the number of sequences from 2,000 to 1,268. Next, we used the criterion of negative design, by only retaining sequences that had above-average Superfamily results. We kept sequences with above-average Superfamily match lengths (above 78) and E-values (log_10_ E *<* −31). This left us with 692 sequences.

Since we planned to test only a few sequences experimentally, we reduced the number of candidates further using four additional criteria: (1) We excluded sequences with below-average similarity scores versus Pfam (the left part of the all-position peak in Fig. 2), leaving 215 sequences. (2) We excluded sequences that had a cavity buried in the predicted 3D structure. (3) We required a total unsigned protein charge of less than 6. (4) We allowed no more than 15 mutations that drastically changed the amino acid type (defined by a Blosum62 similarity score between the two amino acid types of −2 or less). This left us with 16 candidate sequences, shown in Fig. 3. These sequences were separated into four groups, based on visual inspection. Group 2 was eliminated based on its Arg494 residue, absent from CASK homologs. One candidate was selected from each of the other groups (highlighted in Fig. 3), with a preference for native or homologous residue types at positions 492 (candidate 1350), 494 (candidate 1555), and 548 (candidate 1669)— residues that are close to the peptide binding interface. Finally, the three candidates were each simulated by molecular dynamics with explicit solvent for one microsecond, and their stabilities and flexibilities appeared comparable to the wildtype (Supplementary Material, Figs. S1–S2). Therefore, the three sequences were retained for experimental testing. The number of mutations, compared to wildtype CASK, were 50 (candidate 1350), and 51 (candidates 1555 and 1669), representing over 60% of the sequence.

**Figure 3:**
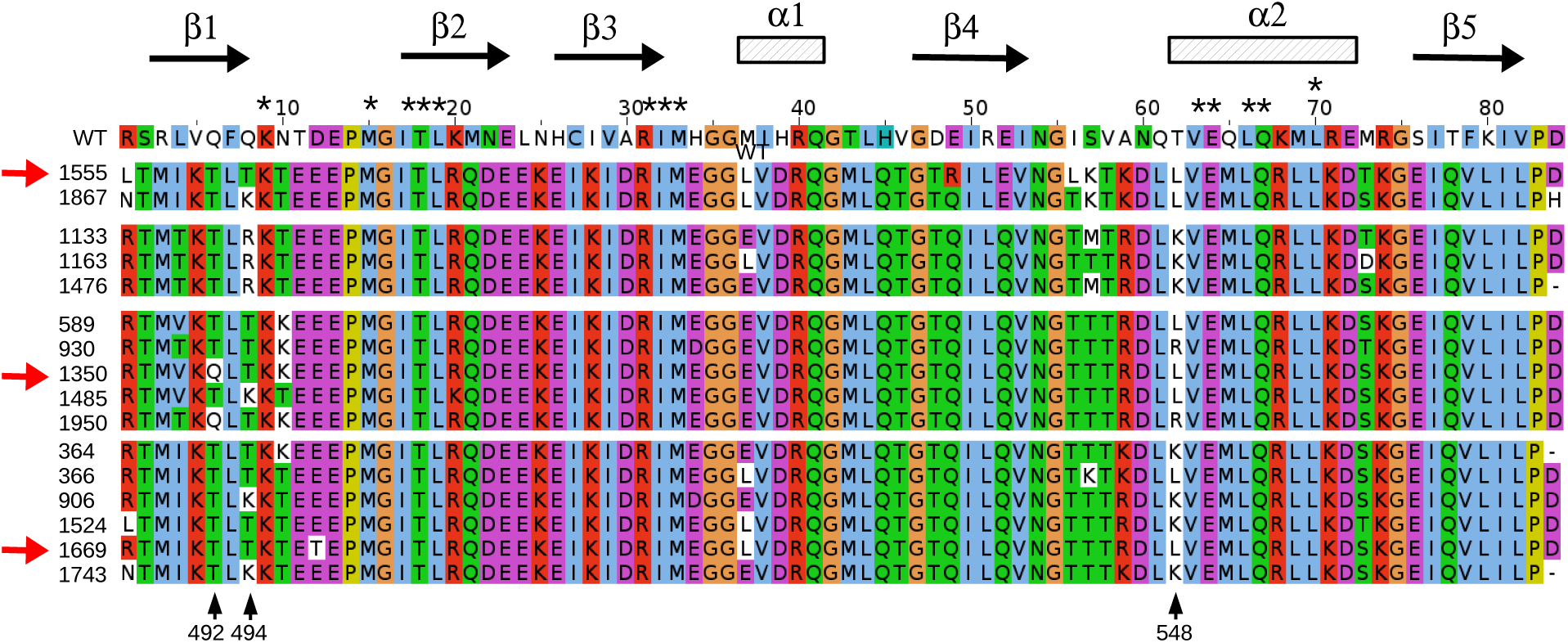
WT and the 16 final candidate designed sequences based on the CASK template. The sequences tested experimentally are indicated by red arrows. Asterisks (above) indicate 13 positions not allowed to mutate during the design, in addition to Gly and Pro residues.

### Experimental characterization of selected sequences

#### Earlier designs based on the Tiam1 template

Computational redesign of Tiam1 was described earlier. ^49^ It used the Tiam1 PDZ domain structure (PDB code 4GVD; Fig. S3) and the simpler, NEA electrostatics model. ^49^ Eight designs were expressed and purified; however, their yields were low. Circular dichroism (CD) yielded spectra typical of random coil polymers, suggesting the proteins were misfolded, whereas the Tiam1 PDZ domain yielded a spectrum typical of a folded structure containing both helical and beta sheet secondary structure (Supplementary Material, Fig. S4). Similarly, 1D-NMR spectra of the amide region of the NEA designs had limited dispersion and broad resonances compared to the native Tiam1 PDZ domain (Fig. S5). Moreover, differential scanning fluorimetry (DSF) in the presence of known Tiam1 ligands did not show any binding by the Tiam1 NEA designs, while the Tiam1 PDZ domain showed robust binding (Suppl. Material, Fig. S6). Together, these data indicate that the NEA-based designs of the Tiam1 PDZ domain could be overexpressed but adopted unfolded structures that lacked the ability to bind known Tiam1 peptide ligands.

#### Designs based on the CASK template

e refer to them as Next, we characterized the three designs selected above, which we refer to as FDB-1350, FDB-1555, and FDB-1669. They were obtained using a new apo CASK PDZ domain structure (PDB code 6NH9) as template and the more rigorous FDB electrostatics model. ^47^ The expression yields in *E. coli* were improved over the NEA Tiam1 designs, though not to the level typically seen with native PDZ domains. In contrast to the NEA Tiam1 designs, CD spectra of the FDB designs were similar to native PDZ domains, suggesting that these designs were structured (Fig 4). 1D-proton NMR of the amide region showed good dispersion and relatively sharp lines, consistent with a folded protein (Fig 5).

**Figure 4:**
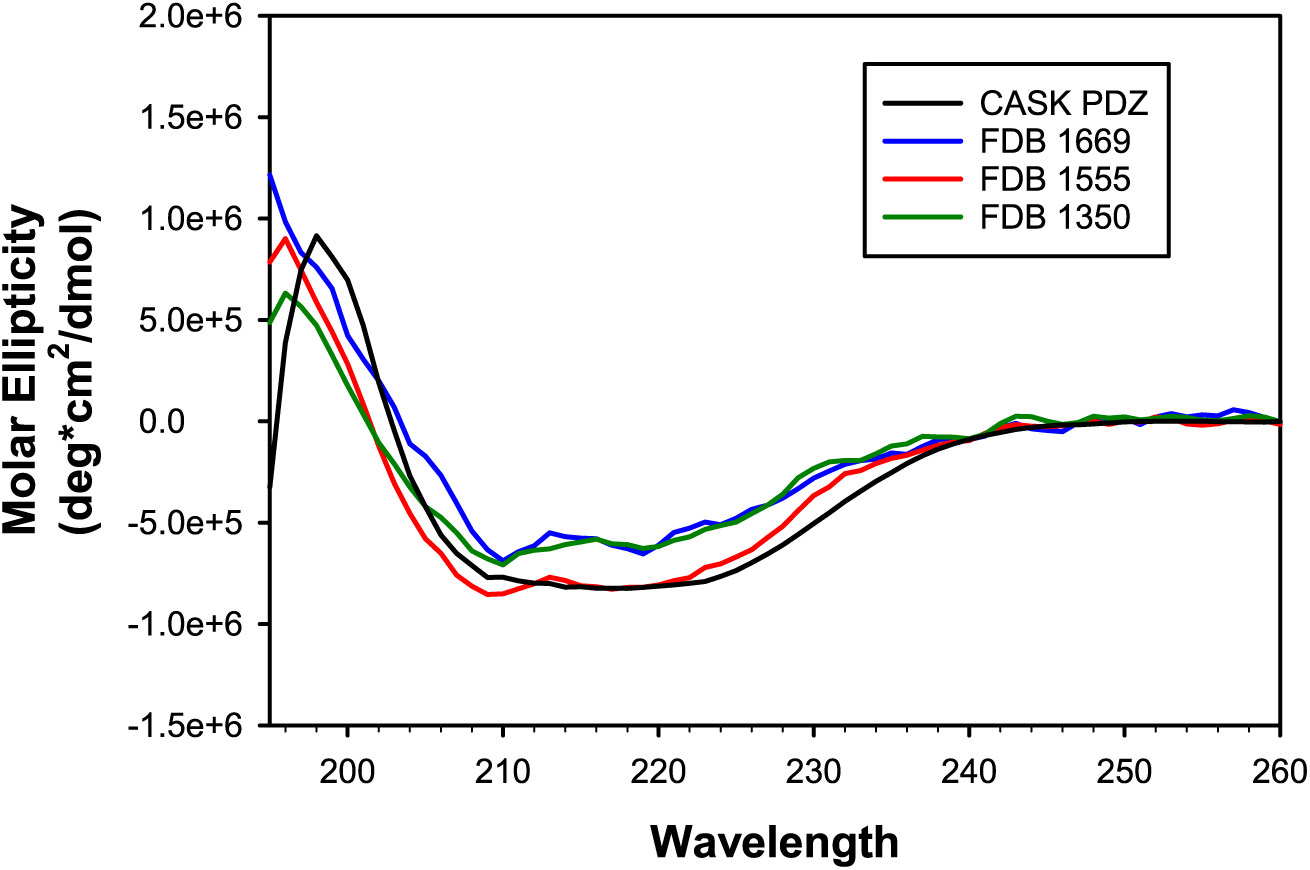
Circular dichroism spectra of a natural PDZ domain (CASK, black) and three selected designs based on the CASK template and the FDB electrostatic model. FDB-1350 (green), FDB1555 (red), and FDB-1669 (blue) all have *α* helix and *β* strand signals similar to a native PDZ domain like CASK (black). The concentration of each protein ranged from 10 to 20 *µ*M.

**Figure 5:**
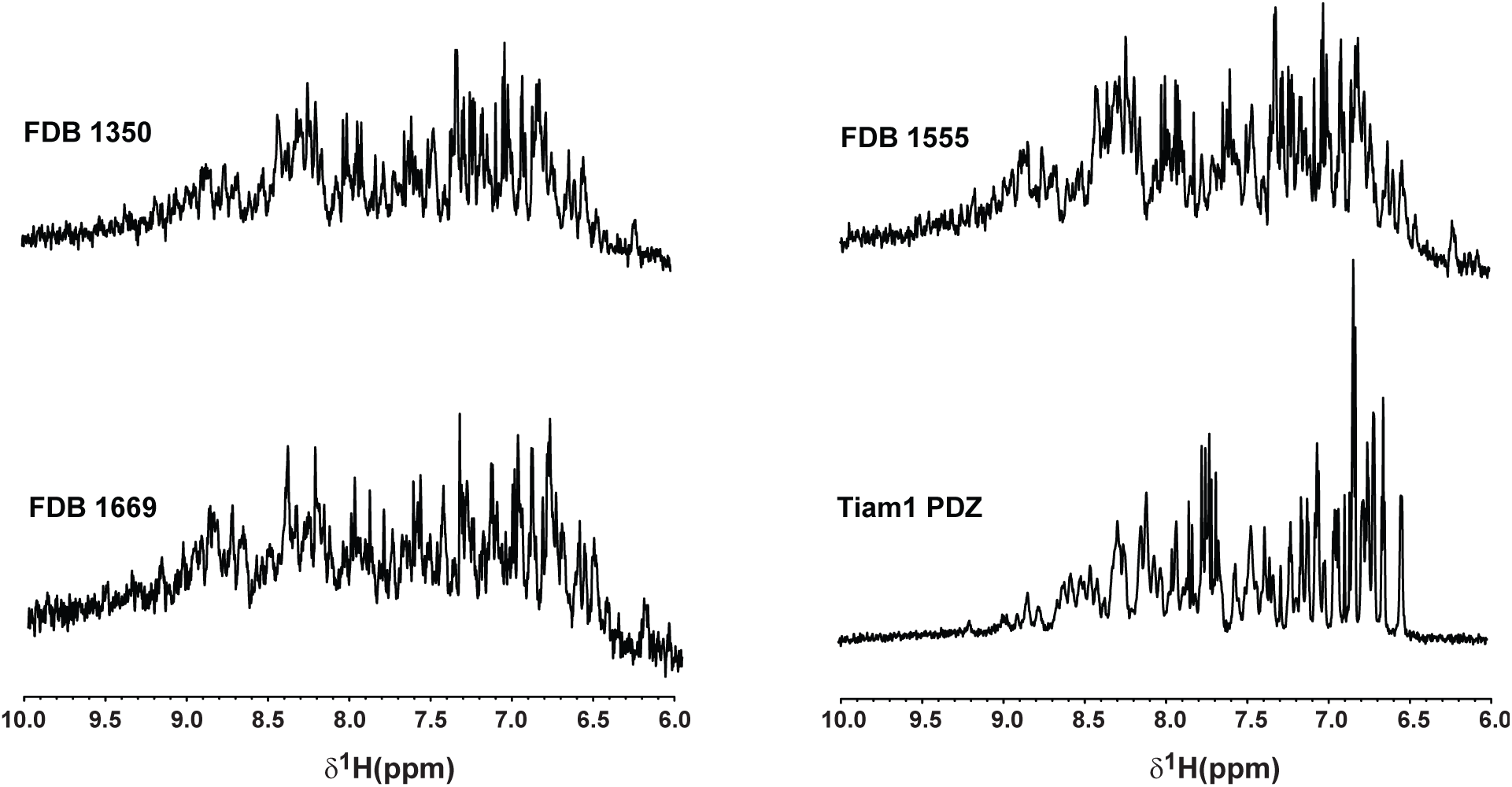
Proton NMR spectra of the natural Tiam1 PDZ domain and three selected designs, obtained based on the CASK template and the FDB electrostatic model.

The design strategy included fixing the sequence of 13 amino acid positions important for peptide binding. Therefore, we tested the ability of the designs to bind CASK ligands, using DSF experiments. The CASK PDZ domain showed binding to SDC1, Caspr4 and NRXN (Fig 6 and Table 1), as expected. Strikingly, two of the three CASK FDB designs characterized also showed binding to some of the peptides. Thus, FDB-1350 had a significant thermal shift in the presence of NRXN and SDC1. FDB-1669 showed a 1.0*^◦^*C change in T_1*/*2_ in the presence of the NRXN peptide. In contrast, FDB-1555 did not show significant thermal shifts in the presence of any peptide. From these data, we conclude that the three CASK FDB designs were folded and two were capable of interacting with peptide ligands.

**Figure 6:**
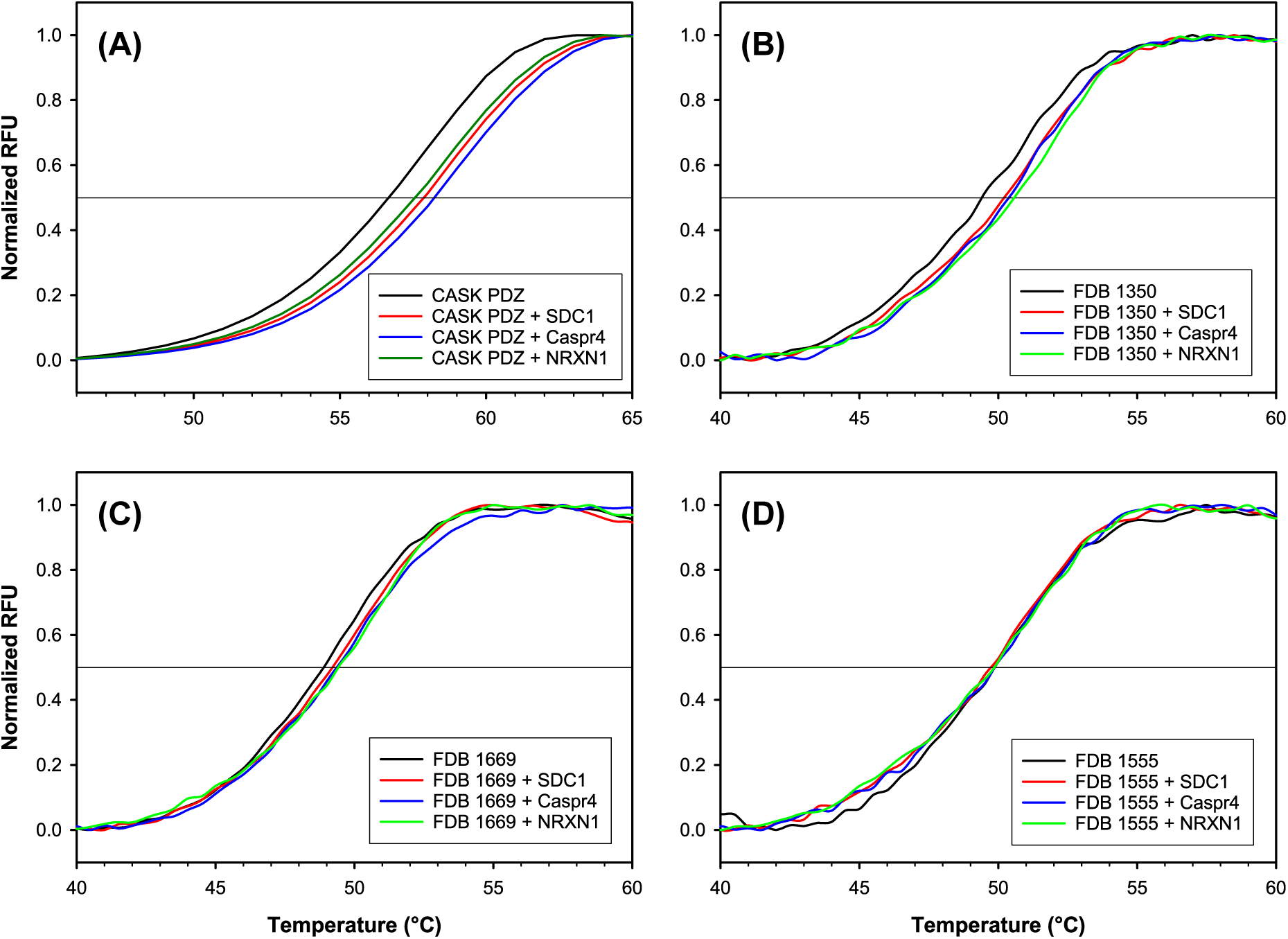
Differential scanning fluorimetry of (A) a natural PDZ domain (CASK) and (B–D) three selected designs based on the CASK template and the FDB electrostatic model. Signals in the absence and presence of the SDC1, Capr4 and NRXN peptides.

## Discussion

Protein folding is usually thought of as being induced by protein–solvent and solvent– solvent interactions,^76, 77^ since folding buries nonpolar groups and allows waters to interact with polar protein groups and other waters. In this picture, the protein dielectric properties play a role, with the low-dielectric interior pushing polar protein groups out towards high-dielectric solvent.^78, 79^ The protein nonpolar surface also plays a role, with exposed surface leading to fewer water–water interactions. ^80–82^ Thus, it is common to discuss protein solvation in terms of nonpolar and electrostatic components, and most implicit solvent models rely on this separation. ^25^ Small proteins have been found to fold correctly in MD simulations with both explicit and accurate implicit solvent models. ^28, 83–86^ The inverse folding problem is arguably even more complex, since it explores an enormous space of sequences, albeit with a reduced conformation set. Modeling the solvent is a key step to solve this problem, and a key ingredient of our design procedure.

The first solvation component in our model is nonpolar and uses accessible surface areas and atomic surface tensions. Nonpolar solvation of a large collection of small molecules correlated well with surface area, ^87^ supporting this treatment. The surface tension parametrization was updated recently, compared to our earlier Tiam1 designs. ^88^ Surface interactions in proteins are complex and have a many-body character,^8, 42^ since three or more residues can have surfaces that all overlap. Our model explicitly includes the most important 3-residue effects, while others are accounted for implicitly.^45^

The largest protein solvation effects are electrostatic, and they also have a many-body character. Indeed, a side chain can desolvate an interacting pair, affecting the strength of their interaction. The electrostatic, Generalized Born component of our model captures this effect. However, for previous Tiam1 design calculations, ^49^ we had used an approximation where each protein residue experienced a constant, native-like, dielectric environment. This removed the many-body character of electrostatic solvation (and made the calculations very efficient). However, the Tiam1 designs were shown here to be largely unsuccessful: the proteins could be overexpressed but were only weakly structured. In contrast, preserving the many-body solvation effects was shown recently to give improved accuracy for side chain pK*_a_*’s.^47^ It also led to increased similarity between CPD sequences and natural sequences of several PDZ proteins. ^47^ Therefore, for the present CASK redesign, we applied the newer, many-body FDB model and obtained improved results.

Our calculations used a CASK X-ray structure first reported here, determined at Å resolution. In our design procedure, the protein backbone was held fixed in the X-ray conformation, while side chains mutated and explored rotamers. More precisely, the backbone motions were not ignored but were treated implicitly, through the protein dielectric constant, *∊_P_*. The value used here, *∊_P_* =4, is known to be physically reasonable for proteins. ^89, 90^ Microsecond MD simulations further showed that the tested sequences, FDB-1350, −1555, and −1669, have backbone structures very similar to the wildtype protein, and have native-like flexibilities.

Although our folded state model is arguably non-empirical, our design procedure did include several empirical elements. First, for the unfolded state, we assumed a simple, extended peptide model, ^91^ to which a correction was added that involved type-dependent amino acid chemical potentials.^43^ These were chosen ^49^ to maximize the likelihood of sampling a collection of natural PDZ sequences. This was possible because our method samples sequences according to a known, Boltzmann probability distribution (known up to a constant normalization factor).

Second, we used several filters to choose a handful of sequences for experimental testing. Starting from sequences within 1.5 kcal/mol of the top folding energy, we used the (computed) isoelectric point to reduce the chances of aggregation. We also used negative design, which was not part of the sequence generation, eliminating candidates with below-average Superfamily fold recognition scores. We also eliminated sequences with below-average similarity scores, relative to natural sequences. These two criteria were actually not very stringent, in the sense that the score distributions were very narrow (Fig. 2). At this point, we were left with 215 sequences. We then eliminated sequences whose structural models included large cavities and ones with a large net charge. Finally, we eliminated sequences with more than 15 “drastic” mutations (corresponding to Blosum scores of −2 or less). This last, purely empirical filter left us with 16 sequences. Among these, we chose three that were representative. If some of the filters were omitted, for example if we had selected sequences randomly at the 215-sequence stage, we expect that there might have been more failures, instead of 2–3 successes out of 3 tests.

The three tested proteins could be overexpressed, had sharp 1D-NMR peaks and native-like CD spectra. Two exhibited a shift of their thermal denaturation in the presence of one or two peptides that are known CASK ligands. The expression yields and protein solubilities were lower than for wildtype CASK, so that it was not possible to produce large amounts of pure protein for 2D-NMR or X-ray crystallogrphy. It may be possible to improve this behavior by testing a larger number of designs and/or using an empirical filtering of candidate sequences for solubility.

The present design method, which combines molecular mechanics, continuum electrostatics, and Boltzmann sampling, is an example of “physics-based” CPD. It is striking and encouraging that this approach allows whole protein redesign to be done successfully. Evidently, the solid physical basis of the energy function and its careful parameterization can lead to a good level of success, despite various approximations. We expect that the “physics-based” route will increasingly yield valuable physical insights and should be a valuable complement to knowledge-based CPD and experimental design.

## Supplementary Material

Supplementary Material is available providing (1) results from MD simulations of Tiam1, CASK and the three selected CASK-based designs; (2) experimental characterization of sequences designed using the Tiam1 template and the NEA electrostatics model; (3) statistical data on the CASK X-ray structure refinement.

## Acknowledgements

Some of the calculations were done at the CINES supercomputer center of the French Ministry of Research. NAMD was developed by the Theoretical and Computational Biophysics Group in the Beckman Institute at the University of Illinois at Urbana. We are grateful to Dr. Jay Nix and the staff at beamline 4.2.2 at the Advanced Light Source, Lawrence Berkeley National Laboratory. We thank Lokesh Gakhar (U of Iowa Protein and Crystallography Facility) for helpful advice in crystallgraphy. The X-ray crystallography software used in the project was installed and configured by SBGrid. The Roy J. Carver Charitable Trust is acknowledged for funding of the Carver College of Medicine Medical Nuclear Magnetic Resonance Facility.

## Supplementary Material

Below, we provide information on the stability and flexibility of the selected CASK-based designs in microsecond molecular dynamics (MD) simulations. Next, we report the experimental characterization of PDZ sequences designed using the Tiam1 template structure and the NEA electrostatics model. Finally, we report the crystallographic structure statistics for the apo CASK PDZ domain.

### Stability of the three selected CASK-based designs in MD

As a first test of the three selected sequences, FDB1350, FDB1555, and FDB1669, they were subjected to MD simulations using an explicit solvent environment, for 1000 ns. Wildtype CASK (WT) was also simulated. Convergence of the simulations was good (based on a principal component analysis, not shown). The WT protein was quite stable, with rms deviations from the starting, X-ray structure of 1–1.5 Å (excluding 3–4 residues at each terminus and one very flexible loop, residues 495–502; see Fig. S1). Deviations from its own mean MD structure were similar (Fig. S1). The designed proteins exhibited only slightly larger deviations from the WT X-ray structure (1.2–1.8 Å) and similar, small deviations from their respective mean MD structures, with no visible drift (Fig. S1).

**Figure S1:**
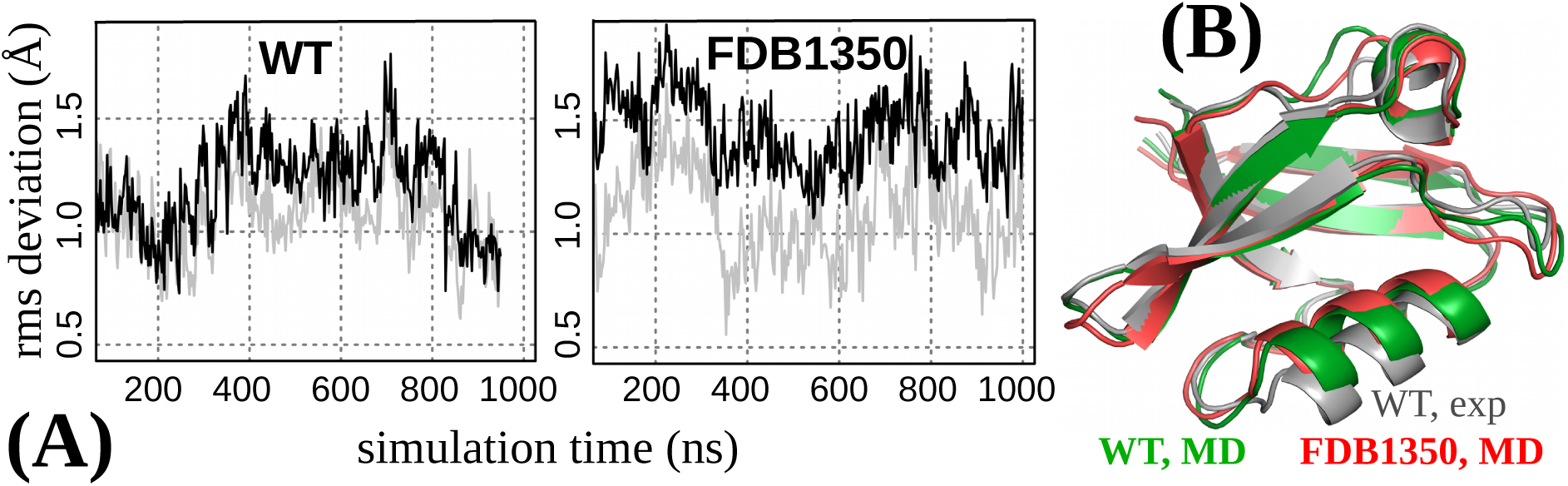
MD simulations of CASK-based designs. **A)** Backbone rms deviations for WT and the FDB1350 designed variant relative to the starting structure (black) and the mean MD structure (grey). **B)** Mean MD structures of WT and designed variant FDB1350.

We also characterized the backbone flexibility of the designed proteins by computing NMR order parameters for the backbone amide groups (Fig. S2). Experimental values were not available for WT CASK, but were available for Tiam1 and a quadruple mutant of Tiam1 (Liu et al, Structure, 12:342, 2016). These proteins were also simulated by MD for one microsecond, with and without the peptide ligands Sdc1 and Caspr4, respectively. In Fig. S2, we show the order parameters for both proteins in the apo and holo states, from experiment (circles) and MD (continuous lines) (top two panels). The agreement is very good. Next, we show (Fig. S2, bottom panel) the order parameters for WT CASK and the three selected CASK-based designs, FDB1350, FDB1555, and FDB1669 (apo proteins). Comparing the designed proteins to WT CASK, the results were similar, with some differences in loop regions. Two designs were slightly less flexible than WT (see positions 492-502 in *β*_1_-*β*_2_, 521-524 in *β*_3_-*α*_1_), while FDB1350 was slightly more flexible (see 492–496 in *β*_1_-*β*_2_ and 559-561 in *α*_2_-*β*_5_). Evidently, the design calculations do not produce overly-rigid or overly-flexible proteins in a systematic way.

**Figure S2:**
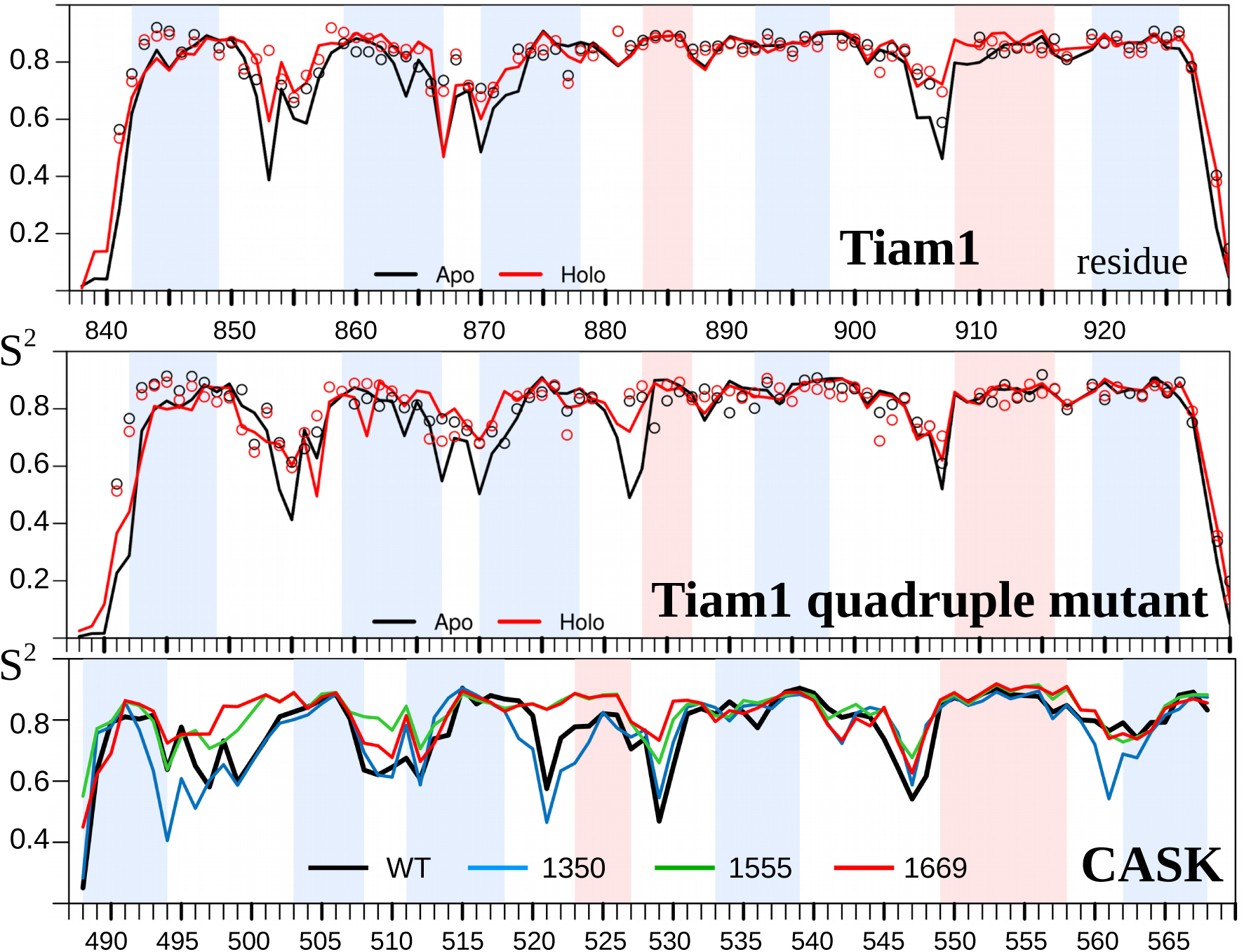
Backbone amide NMR order parameters for natural and designed proteins. **Top panel:** Tiam1 with and without the Sdc1 peptide ligand. Circles are experimental values; lines are from *µ*sec MD simulations. **Middle panel:** analogous data for the Tiam1 quadruple mutant and the Caspr4 peptide. **Bottom panel:** Apo WT CASK and the three designed variants; values from MD.

### Experimental characterization of Proteus designs obtained with the Tiam1 template and the NEA electrostatic model

**Figure S3:**
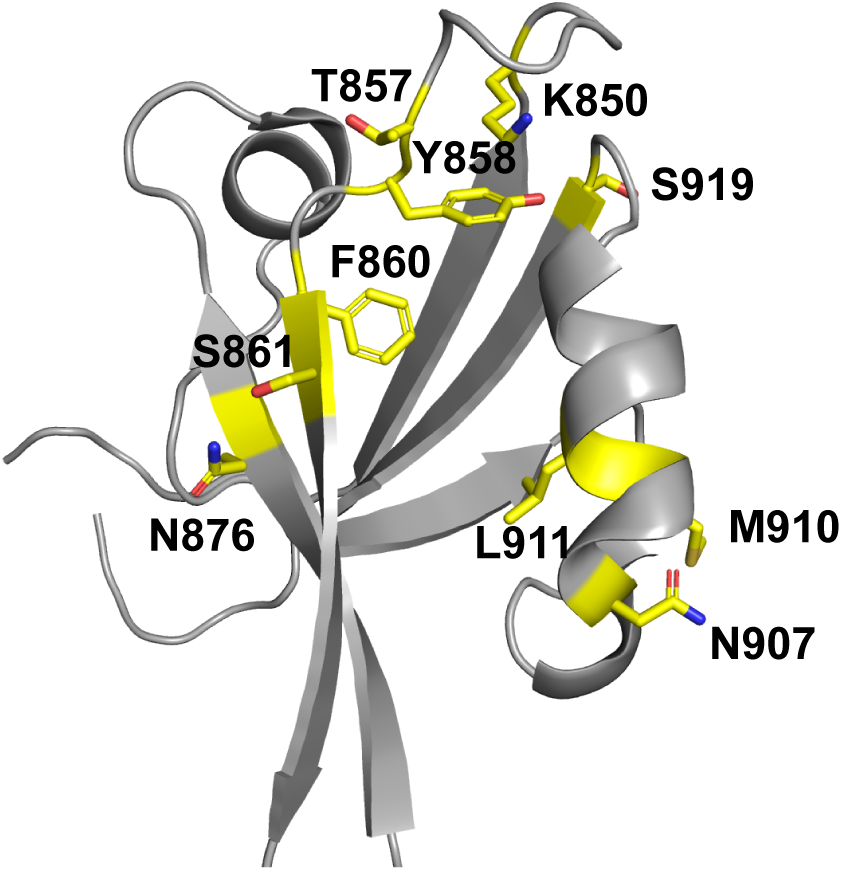
Tiam1 structure. Yellow: 13 positions whose types were fixed in the Proteus designs.

**Figure S4:**
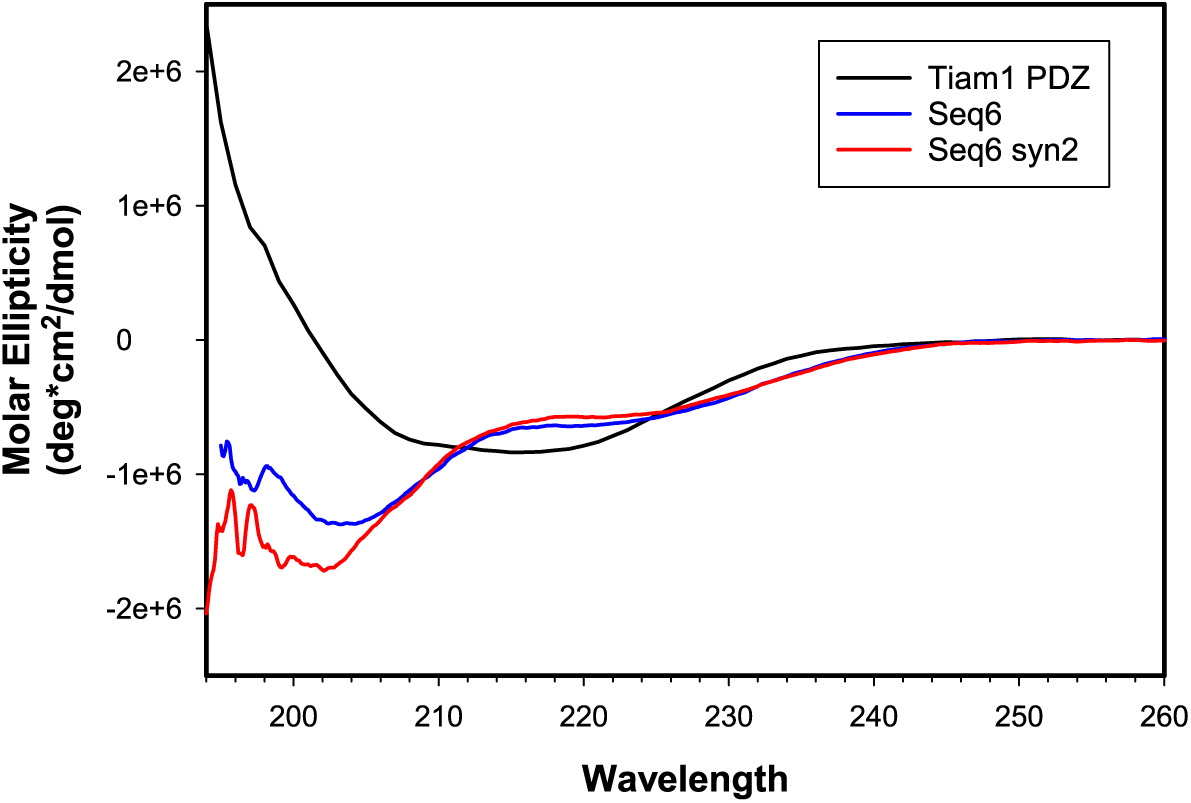
CD spectra of Tiam1 and two designs based on the Tiam1 template and NEA electrostatics.

**Figure S5:**
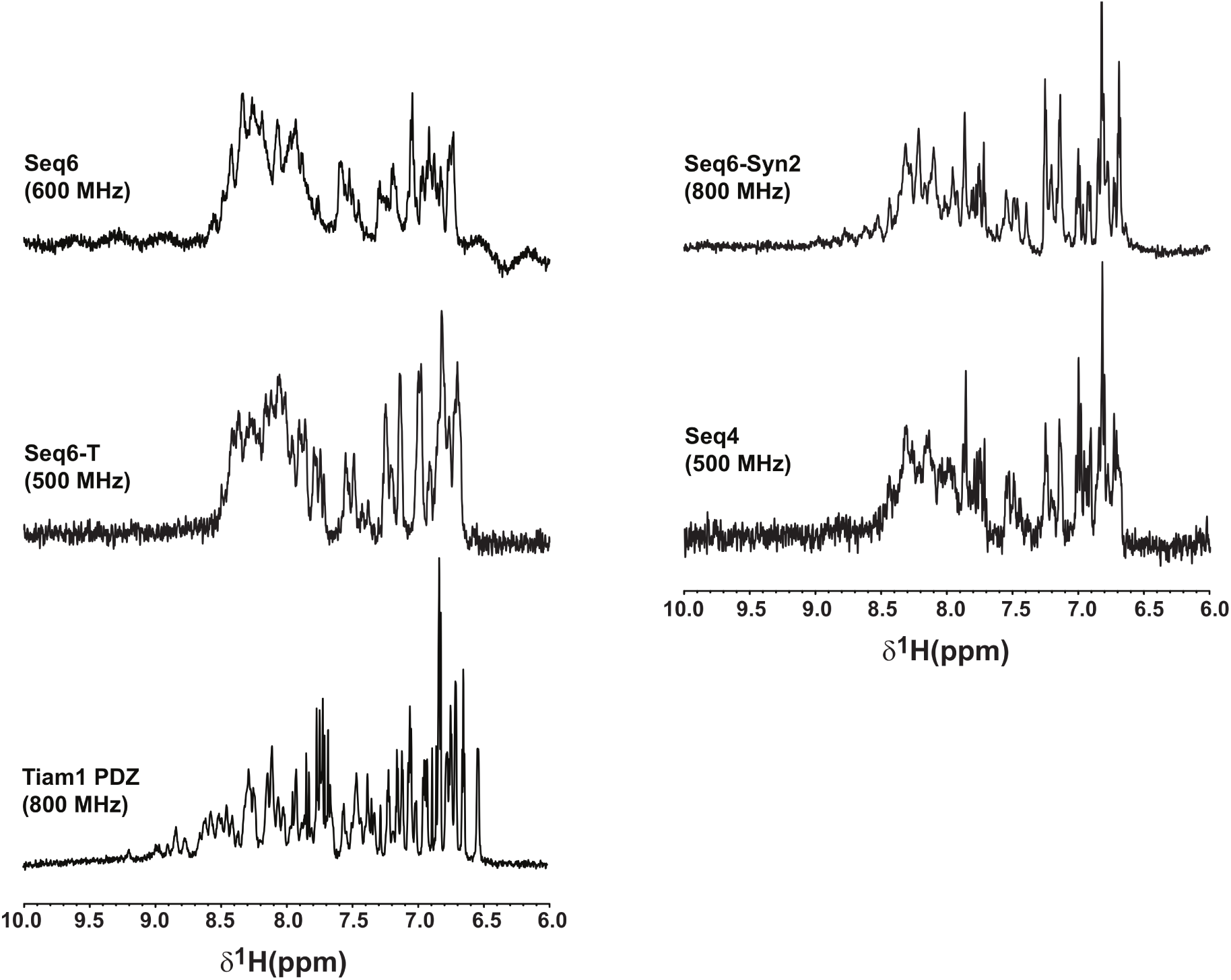
Proton NMR spectra of the Tiam1 PDZ domain and four designs obtained with the Tiam1 template and NEA electrostatics.

**Figure S6:**
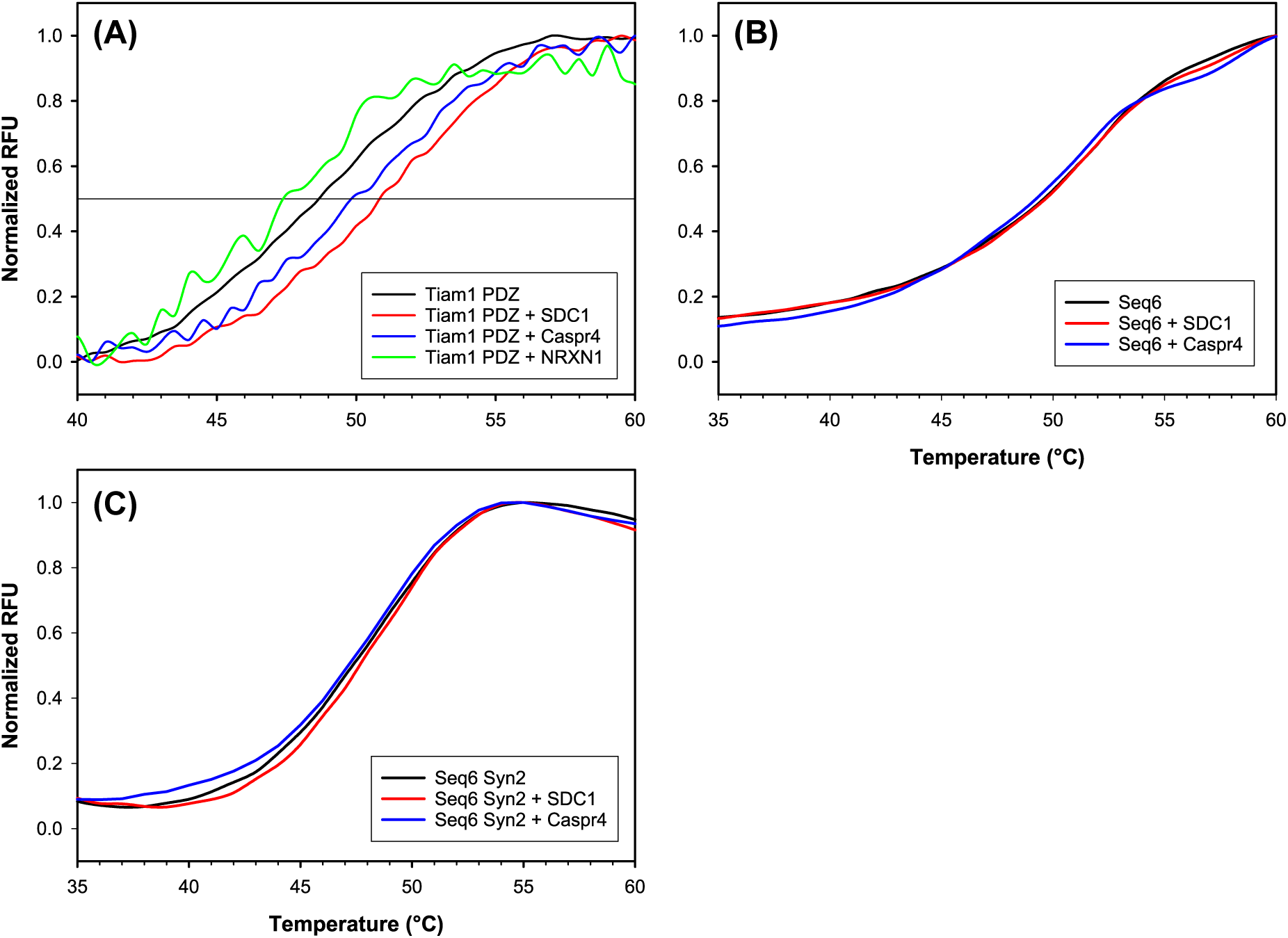
Di_erential scanning uorimetry of a natural PDZ domain (Tiam1) and two designs based on the Tiam1 template and the NEA electrostatic model. Signals in the absence and presence of the SDC1, Capr4 and NRXN peptides.

### Human apo CASK PDZ domain X-ray structure statistics

**Table S1:** Crystallographic statistics for the human apo CASK PDZ domain

| Data collection statistics |  |
| --- | --- |
| Beam line | ALS 4.2.2 |
| Wavelength (Å) | 1.0003 |
| Space group | C 1 2 1 |
| Unit cell dimensions (a, b, c) (Å) | 61.1, 35.4, 119.5 |
| Unit cell dimensions ($\alpha$, $\beta$, $\gamma$) | 90°, 90.3°, 90° |
| Resolution range (Å) | 59.8—1.85 |
| Total reflections | 37,385 (7,461) |
| Unique reflections | 20,769 (1,910) |
| Multiplicity | 1.8 (1.7) |
| Completeness (%) | 93.7 (93.7) |
| I/$\sigma$ (I) | 10.4 (2.1) |
| Wilson B-factor (Å<sup>2</sup>) | 50.7 |
| R<sub>meas</sub> | 0.030 (0.402) |
| CC<sub>1/2</sub> | 99.8 (91.1) |
| Refinement statistics |  |
| Resolution (Å) | 1.85 |
| No. of reflections used in refinement | 20,739 (2,705) |
| No. of reflections used for R<sub>free</sub> | 964 (133) |
| R<sub>work</sub>/R<sub>free</sub> | 0.226/0.263 |
| No. of atoms | 4,188 |
| Protein | 4,037 |
| Water | 151 |
| B-factors (Å<sup>2</sup>) | 53.0 |
| R.M.S.D.<sup>a</sup> |  |
| Bond length (Å) | 0.29 |
| Bond angle (degrees) | 0.46 |
| Ramachandran plot statistics (%) |  |
| In preferred regions | 98.0 |
| In allowed regions | 2.0 |
| Outliers | 0.0 |
| PDB accession code | 6NH9 |

The numbers in parentheses are for the highest-resolution shell. <sup>a</sup>Root mean square deviation from ideal values.

